# Association of GADD45A and favorable outcome in hormone positive breast cancer

**DOI:** 10.1101/2025.06.02.657315

**Authors:** Chih-Yi Lin, Chun-Yu Liu, Ta-Chung Chao, Chi-Cheng Huang, Yi-Fang Tsai, Ling-Ming Tseng, Jiun-I Lai

**Author notes:** Correspondence: Jiun-I Lai.

## Abstract

**Background:** Hormone receptor-positive (HR(+)) breast cancer is the most prevalent subtype in breast cancer, and although treatment options are rapidly increasing, still poses as an unmet need in advanced cases. Growth arrest and DNA damage-inducible alpha (GADD45A) is a gene with known relationship with the cell cycle and DNA damage pathways. Conflicting roles in breast cancer biology have been reported for GADD45A. In our study, we investigate the clinical and molecular association of GADD45A in HR(+) breast cancer, evaluating its prognostic potential and relationship with estrogen signaling.

**Methods:** GADD45A expression was assessed in 100 breast cancer tissue samples from a tissue microarray and correlated with clinicopathologic parameters. Public genomic datasets were analyzed to validate associations between GADD45A expression and clinical outcomes in HR(+)HER2(−) breast cancer. Gene set enrichment analysis (GSEA) was used to investigate pathway correlations.

**Results:** GADD45A protein levels were highest in HR(+)HER2(-) tumors and positively correlated with estrogen receptor (ER) levels, consistent with a luminal phenotype. No significant association was observed between GADD45A levels and progression-free survival in CDK4/6 inhibitor-treated patients, suggesting limited predictive utility in this context. However, higher GADD45A expression was significantly associated with improved overall and relapse-free survival in HR(+)HER2(-) breast cancer in the METABRIC cohort, but not in HER2(+) or triple- negative subtypes. GSEA revealed a positive association between GADD45A expression and the estrogen late response signature, suggesting a functional link with hormone signaling.

**Conclusion:** GADD45A expression is enriched in luminal-type HR(+) breast cancer and correlates with favorable clinical outcomes. While not predictive of CDK4/6 inhibitor response, GADD45A may serve as a surrogate marker of estrogen signaling and improved prognosis. These findings support further investigation into GADD45A as a biomarker for stratification and potentially as a therapeutic target in HR(+) breast cancer.

## Introduction

Breast cancer remains the most frequently diagnosed cancer and a leading cause of cancer mortality among women worldwide ^1^. Breast cancer can be classified according to molecular subtypes into hormone receptor-positive (HR(+), HER2(-)), HER2 (HER2(+)) and triple-negative breast cancer (TNBC) ^2^. HR+/HER2− breast cancer accounts for the majority of cases, ranging from approximately 60% to 70% globally ^3, 4^. In HR(+) breast cancer, CDK4/6 inhibitors combined with endocrine therapy is the standard of care in metastatic and early disease ^5^. Newer developments in advanced HR(+) breast cancer include elacestrant, a selective estrogen receptor degrader, showing efficacy in ESR1-mutated tumors ^6^, trastuzumab-deruxtecan showing unprecedented activity in HER2-low metastatic breast cancer, and capivasertib acquiring FDA approval for PIK3CA/AKT1/PTEN-altered tumors ^7^, among others. However, despite much progress in drug development, resistance ultimately occurs that lead to treatment failure, and metastatic HR(+)HER2(-) breast cancer currently still remain incurable, highlighting need for novel insights and innovation.

In our previous study, we discovered that histone deacetylase inhibitors upregulate Hippo pathway downstream genes, including GADD45A, and we reported that GADD45A was associated with survival benefit in HR(+) breast cancer patients ^8^. GADD45A is a member of the GADD45 family of stress sensors, activated by genotoxic and oncogenic stress, is involved in maintaining genomic stability and regulating cellular damage response^9^, and has been linked to tumorigenesis, apoptosis, and tumor progression ^10^. GADD45A can function as tumor suppressing or as tumor promoting, depending on the biological context. In ras-driven breast tumorigenesis, GADD45A acts as a tumor suppressor by enhancing JNK-mediated apoptosis and p38- mediated senescence^11^. Conversely, in Myc-driven breast cancer, GADD45A promotes tumor growth by modulating GSK3β/β-catenin signaling and reducing MMP10 expression, thereby increasing tumor vascularization and progression ^11^. Experimental studies reveal that GADD45A inhibits cancer cell migration and invasion by regulating genes involved in cell adhesion, extracellular matrix (ECM) interaction, and focal adhesion pathways. Loss of GADD45A leads to increased cell motility and invasive potential, partly through altered expression of matrix metalloproteinases and ECM components ^12^. In HR+ breast cancer, GADD45A has been implicated in modulating hormone signaling and cellular stress responses^13^. It is a downstream target of p53 and BRCA1, both of which are critical in HR+ tumor biology ^13^. GADD45A can influence cell cycle arrest and apoptosis in response to DNA damage, potentially affecting the sensitivity of HR+ tumors to endocrine therapies and chemotherapy. Given its involvement in DNA repair and cell cycle regulation, GADD45A may contribute to therapy resistance in HR+ breast cancer. Its modulation could represent a therapeutic target, especially in tumors with defective p53 or BRCA1 pathways ^11^.

Although previous studies have shown GADD45A related to In breast cancer, the association between GADD45A and survival regarding different subtyptes are yet to be determined. In our study, we aimed to investigate the levels of GADD45A in patient samples and from clinical-genomic databases.

## Materials and methods

### Immunohistochemistry (IHC)

Sections from tumors of 17 TVGH patients or tissue microarrays (TMAs) were deparaffinized in UltraClear (J.T.Baker #UN3259) and rehydrated through a descending graded series of alcohols then subjected to antigen retrieval. IHC was performed using the Novolink Polymer Detection System (Leica Biosystems) according to the manufacturer’s instructions. The BC081116e (TissueArray.Com) tissue microarray comprises breast cancer samples with adjacent normal breast tissue, including invasive carcinoma of no special type, breast carcinoma with apocrine differentiation, and adjacent tissue. Pathological grading, IHC data for ER, PR, HER2, and Ki-67, as well as TNM staging according to the AJCC 7th edition, are provided. The array includes 107 cases represented by 110 cores, each with a diameter of 1.5 mm.

### cBioPortal analysis

We queried cBioPortal for Cancer Genomics (http://cbioportal.org) for analyzing complex and multifactorial cancer genomic data from METABRIC ^14^, that included gene expression level, age at diagnosis, ER, PR and HER2 status, OS, RFS and inferred menopausal in “METABRIC Breast cancer cohort (1894 cases in Breast Invasive Ductal Carcinoma, Breast Mixed Ductal and Lobular Carcinoma and Breast Invasive Lobular Carcinoma)”. The OncoPrint, survival tabs and other web-based analysis were applied pursuant to cBioPortal instructions.

### Gene set enrichment analysis (GSEA) and functional enrichment analysis

Gene set enrichment analysis (GSEA) was performed using the GSEA software. Significantly enriched signaling pathways were identified based on normalized enrichment scores (NES) with a false discovery rate (FDR) threshold of <0.05. The analysis was performed by R, using the packages “clusterProfiler”, “tidyverse”, “edgeR”.

## Results

### 1. GADD45A protein levels are associated with estrogen receptor positivity and lower Ki-67 levels in HR(+) HER2(-) breast cancer samples

To quantify GADD45A protein levels in patient tumor samples, we performed immunohistochemistry on tissue microarray slides from a total of 100 breast cancer samples. The clinical characteristics for the tissue microarray cohort is listed as table 1. In the samples analyzed, we observed predominantly nuclear GADD45A immunoreactivity. Staining intensity was scored as negative (−), weak (+), or strong (++) (**Figure 1A**). Patients were stratified by stage, with stage II comprising ∼70% and stage III ∼30%. Positive staining for GADD45A (including intensities 1+ and 2+, denoted as “+” and “++”, respectively) accounted for 59.7% of stage II and 66.6% of stage III cases. No significant difference in GADD45A staining intensity was observed between stages (**Figure 1B**). When categorized by nuclear grade, Grade 1, 2 and 3 each consisted of 3.1%, 63.9%, and 33% respectively. We observed positive GADD45A staining (staining intensity 1+ and 2+ combined) in 100% of Grade 1, 58% of Grade 2, and 62.5% of Grade 3 (note: only 3 samples were grade 1) (**Figure 1C**).

**Figure 1.**
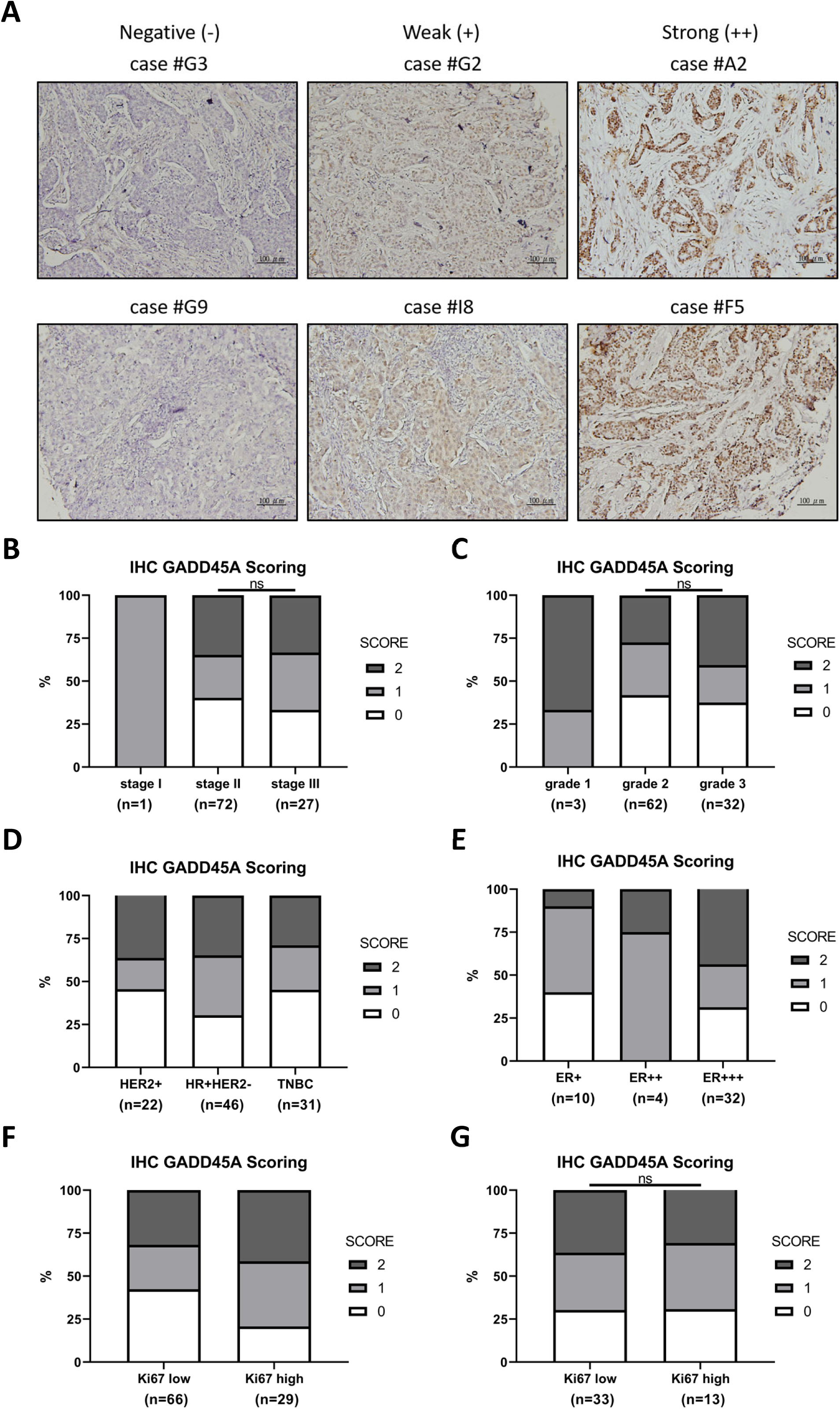
Representative images of tissue microarray (TMA) of GADD45A protein in breast cancer. (**A**), Representative cores of GADD45A protein expression from the TMAs graded on a scale from 0 to 2+ for protein staining intensity. (**B-G**), IHC staining intensity was quantified using stacked bar graphs based on categorical grouping criteria.“ns” indicates not significant (p>0.05).

**Table 1.**
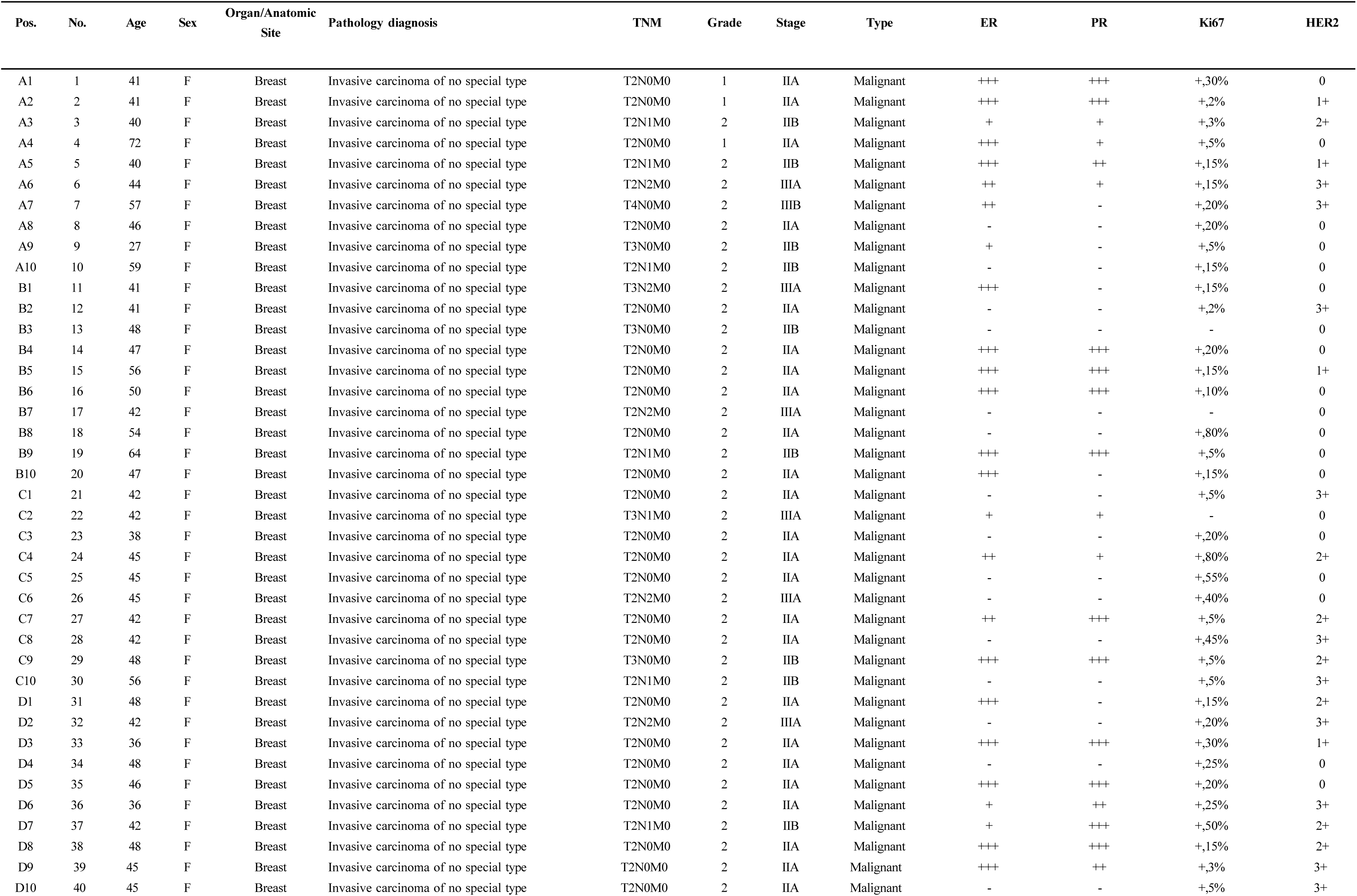
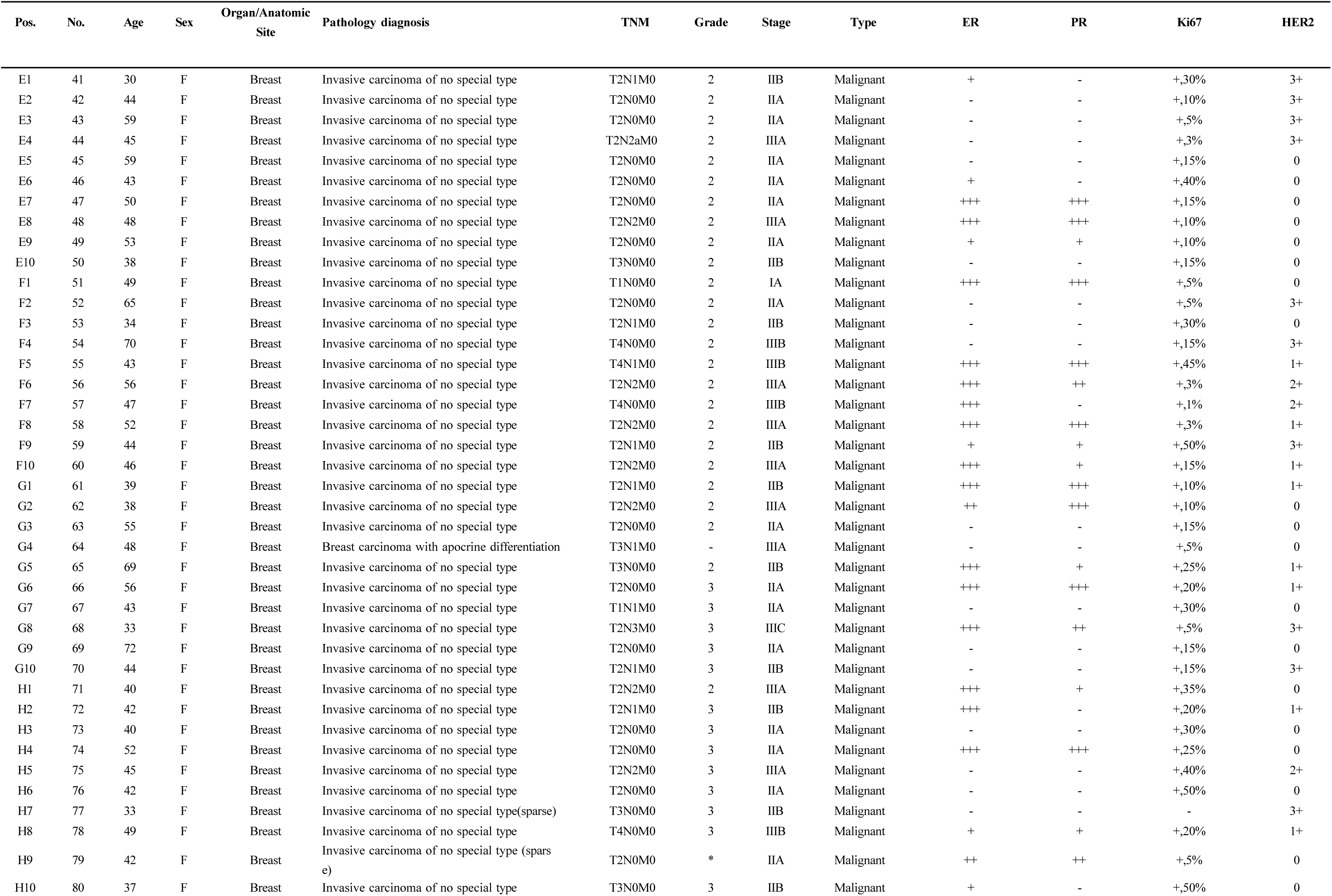
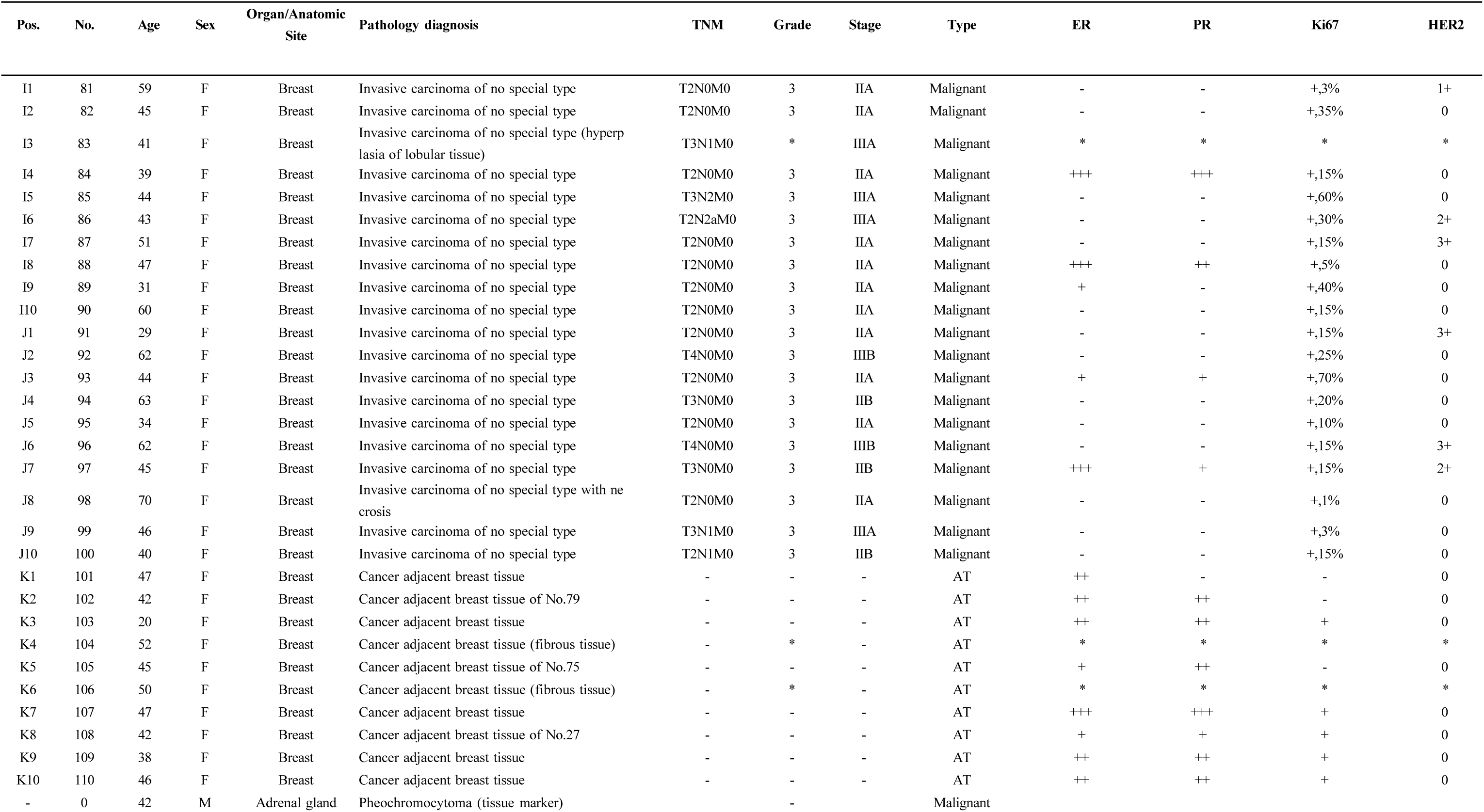
Clinical characteristics for the tissue microarray cohort BC081116e.

We then asked whether there was an association between GADD45A staining intensity and breast cancer molecular subtype. The cohort comprised of 22% HER2(+), 46% HR(+), and 31% TNBC cases, with HR(+) being the predominant subtype. We observed positive staining in 54.6% of HER2 (+), 69.6% of HR(+), and 54.8% of TNBC (**Figure 1D**). Our results reveal that positive GADD45A staining was most prevalent in HR(+) subtype. We then asked whether there was a correlation between ER positivity (graded as +, ++, +++ in this cohort). Interestingly, we observed a positive correlation between ER expression levels and GADD45A strong staining (2+) status, increasing from 10% to 25% and 43.8% across low to high ER groups (**Figure 1E**).

The MONARCHe study highlighted the prognostic value of a 20% cut-off for Ki-67 for recurrence in high risk HR(+)HER2(-) early breast cancer, regardless of whether CDK4/6 inhibitors were used^16^. Therefore, we examined the correlation between high and low Ki-67 (using 20% cutoff) and GADD45A levels. We observed that GADD45A positivity was higher in the patients with higher Ki-67 compared to lower (79.3% versus 57.6%) in allcomers. (**Figure 1F**). However, when we only examined HR(+) patients, we observed that the positivity of GADD45A was 69.2% in high and 69.7% in low Ki- 67 samples (**Figure 1G**), that was no significant difference.

In summary, from GADD45A expression patterns in tissue microarray studies, we observed that HR(+) subtype had the highest prevalence of GADD45A positivity (70%) compared to HER2(+) and TNBC, and higher GADD45A positivity was associated with ER positivity as well as lower Ki-67 levels. Our findings indicate that higher GADD45A is associated with a more luminal phenotype in breast cancer.

### 2. GADD45A levels are not associated with CDK4/6 treatment outcomes

Since GADD45A was highly associated with HR(+) breast cancer, we specifically examined its role in the context of luminal type disease. CDK4/6 inhibitors are currently the standard of care in HR(+)HER2(-) metastatic breast cancer patients, as well as in high risk, resected HR(+)HER2(-) early breast cancer patients. We wished to investigate whether outcome on CDK4/6 inhibitors treatment was associated with GADD45A levels. We collected tumor samples from 16 patients at our institute. These patients were HR(+) HER2(-) metastatic breast cancer patients, all treated with CDK4/6 inhibitors plus endocrine therapy for first line treatment (clinical characteristics in **table 2**). The median age of diagnosis was 61.5, all were postmenopausal, and all received either palbociclib (6 patients) or ribociclib (10 patients) as CDK4/6 inhibitor treatment. 9 patients (56.3%) had progressed on CDK4/6 inhibitors, and the median progression free survival (PFS) was 348 days in these patients. When we divided the patients according to GADD45A staining of their tumor samples, we observed that there was no statistically significant difference between PFS on CDK4/6 inhibitors and GADD45A levels (**Figure 2A-C**). A recent study ^17^ deposited patient tumor baseline RNA-seq information in GEO as well as clinical outcome on palbociclib. (accession GSE186901). In 64 patients with available GADD45A information before starting palbociclib treatment, we also did not observe an association between GADD45A and CDK4/6 inhibitor treatment outcome (**Figure 2D**).

**Figure 2.**
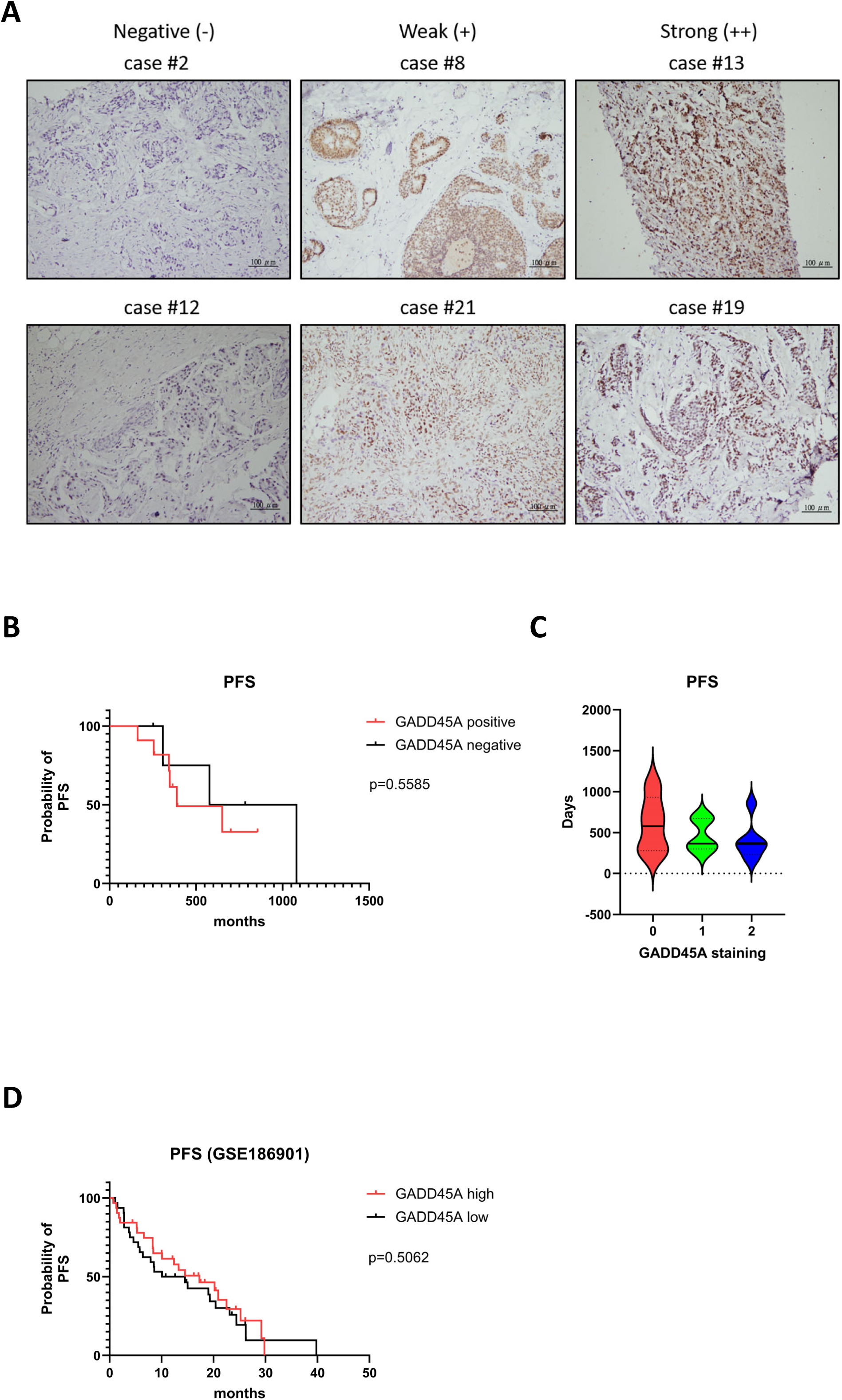
Representative images of GADD45A protein from a clinical cohort. (**A, C**), Representative cores of GADD45A protein expression from the TVGH breast cancer patients graded on a scale from 0 to 2+ for protein staining intensity. (**B**), PFS according to high or low GADD45A levels with HR+ breast cancer patients. (**D**), PFS according to high or low GADD45A levels from RNA-seq information in GEO (GSE186901).

**Table 2.**
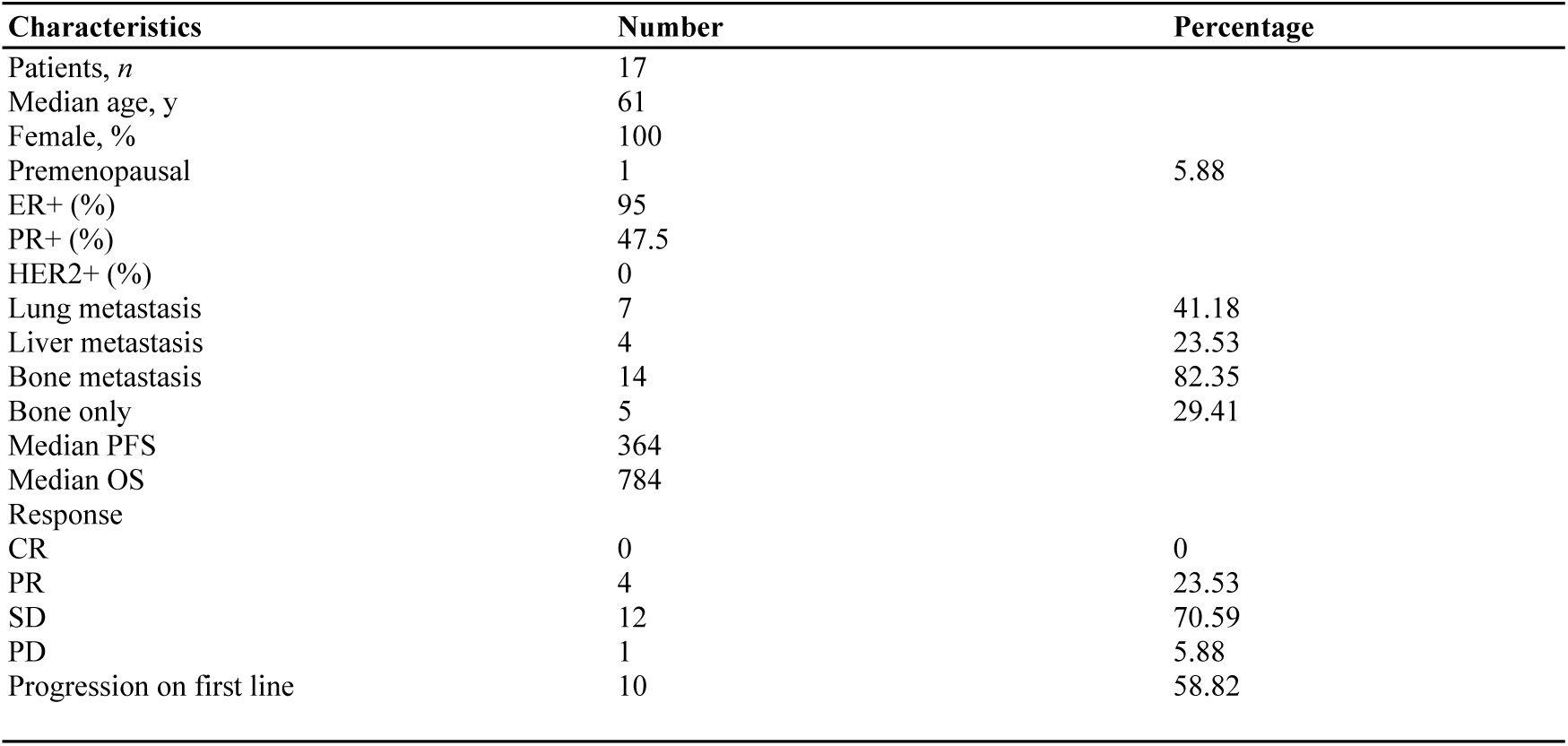
Clinical characteristics of patients at Taipei Veterans General Hospital.

### 3. GADD45A expression is associated with improved outcomes in HR(+) breast cancer

We then investigated the landscape of GADD45A expression in METABRIC database. From the METABRIC database, the number of samples with analyzable GADD45A gene expression levels were 236, 1342, 304 for HER2(+), HR(+)HER2(-), and TNBC subtypes, respectively. We had previously reported that in the complete cohort, high GADD45A was associated with significant overall survival benefit ^8^. When RFS was examined, a similar trend was observed with higher GADD45A associated with longer RFS (**Figure 3A**) compared to low expression. When focusing on the HR(+) HER2(-) subgroup, patients with high GADD45A expression exhibited significantly improved OS (**Figure 3B**) and RFS (**Figure 3C**) compared to those with low expression. In the HER2(+) subtype, either in total, or either HER(+) HR(+) or HER2(+)HR(-), there were no association between GADD45A gene expression and OS (**Figure 3D-F**) or RFS (**Figure 3G-I**). Similarly, TNBC patients showed no correlation with GADD45A and OS or RFS (**Figure 3J-K**). Interestingly, we observed that in the overall population, there were higher GADD45A levels in premenopausal patients compared to postmenopausal patients (**Figure 3L**). This phenomenon was consistent when only HR(+) HER2(-) patients were examined (**Figure 3M**).

**Figure 3.**
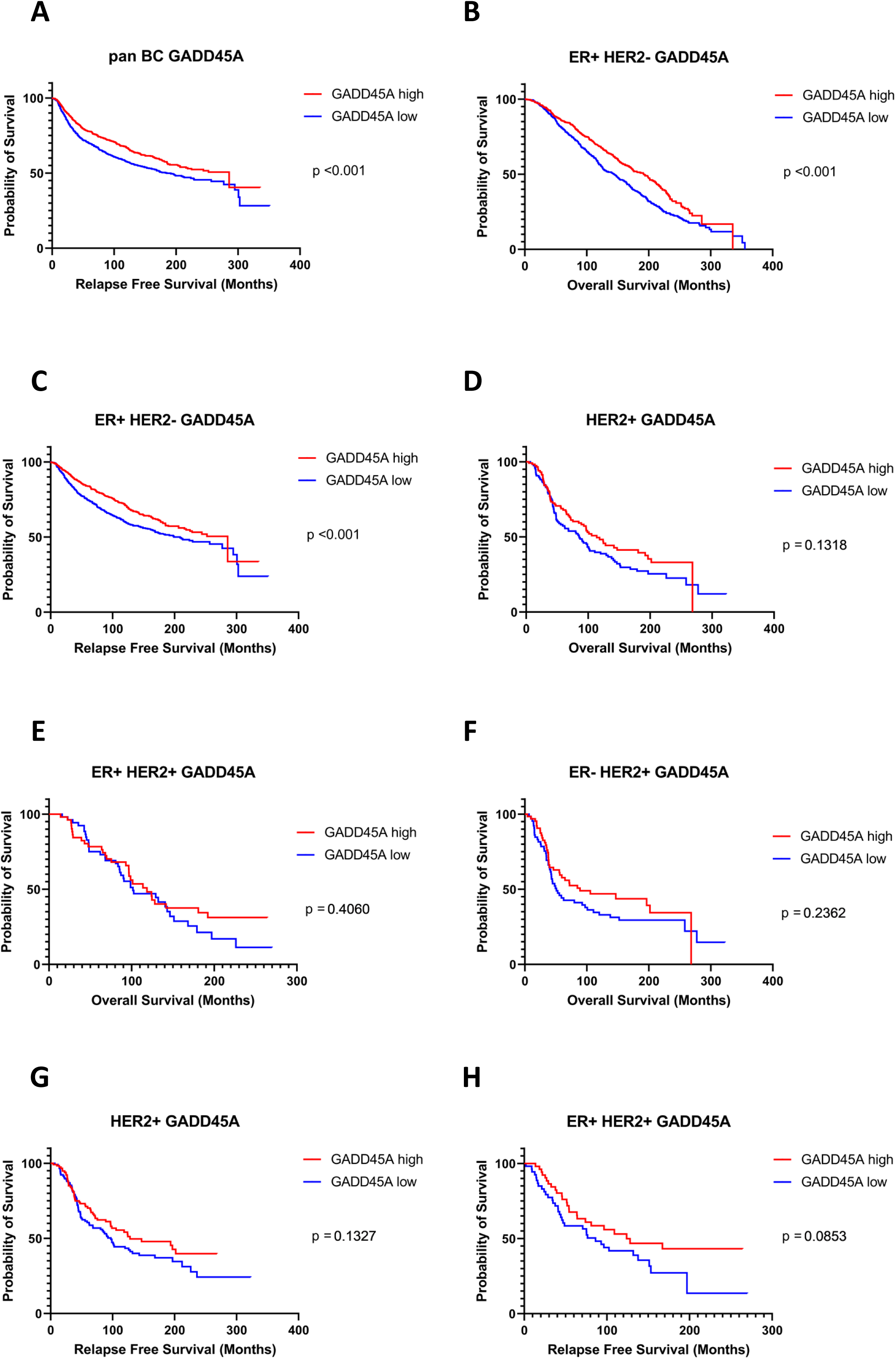
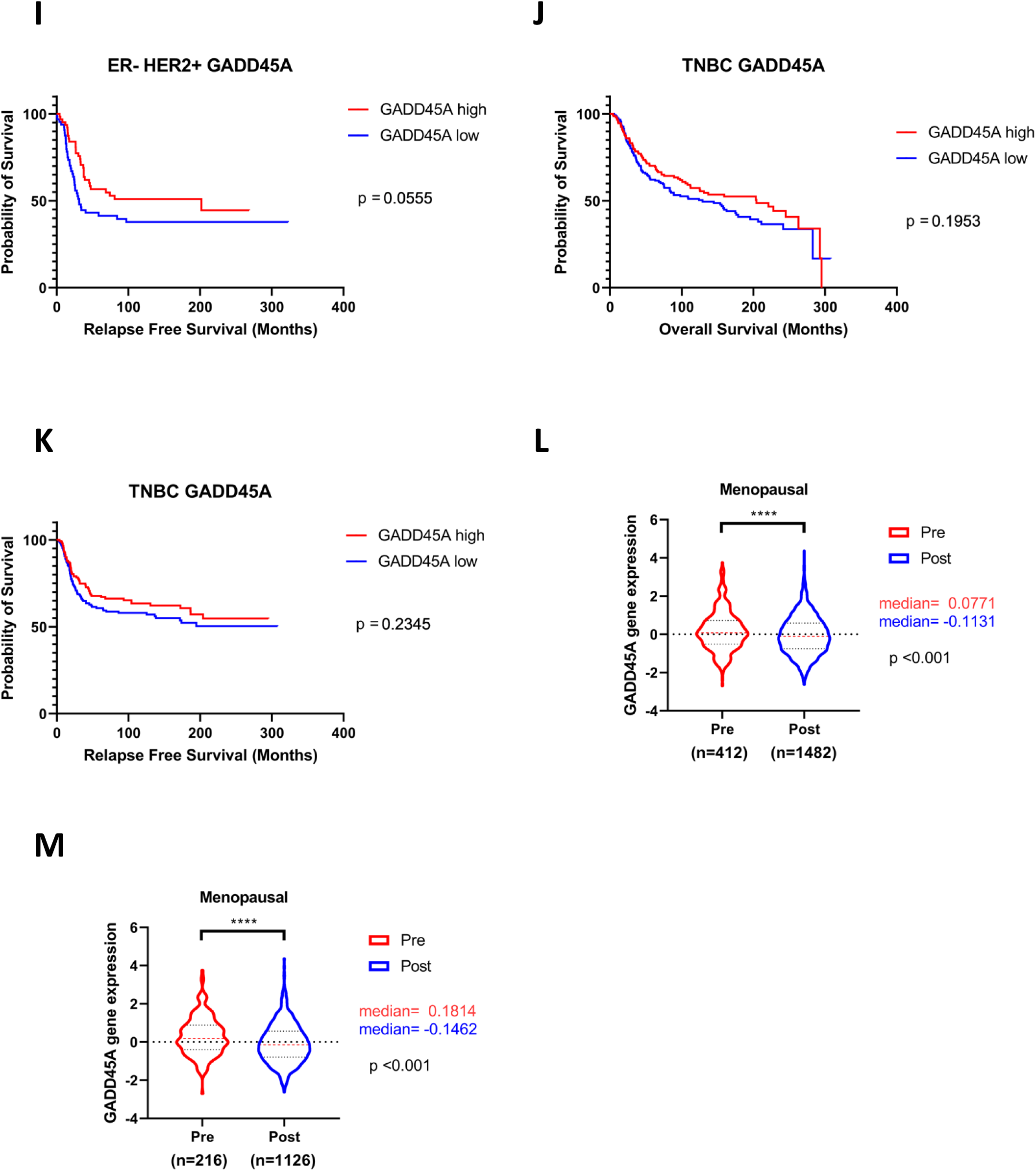
High GADD45A expression significant correlate with improved OS, RFS, and premenopausal status in HR+ BC cohorts. (**A**), Relapse free survival status according to GADD45A in all breast cancer subtypes. (**B**), Overall survival status according to GADD45A in ER+ HER2- breast cancer. (**C**), Relapse free survival status according to GADD45A in ER+ HER2- breast cancer. (**D**), Overall survival status according to GADD45A in HER2+ breast cancer. (**E**), Overall survival status according to GADD45A in ER+ HER2+ breast cancer. (**F**), Overall survival status according to GADD45A in ER- HER2+ breast cancer. (**G**), Relapse free survival status according to GADD45A in HER2+ breast cancer. (**H**), Relapse free survival status according to GADD45A in ER+ HER2+ breast cancer. (**I**), Relapse free survival status according to GADD45A in ER- HER2+ breast cancer. (**J**), Overall survival status according to GADD45A in TNBC breast cancer. (**K**), Relapse free survival status according to GADD45A in TNBC breast cancer. (**L**), The association between GADD45A expression levels and menopausal status was analyzed using violin plots in all breast cancer subtypes. (**M**),The association between GADD45A expression levels and menopausal status was analyzed using violin plots in ER+ HER2- breast cancer. ****p<0.001.

### 4. GADD45A expression is associated with estrogen signaling pathways

Since our above data revealed that GADD45A was associated with hormone positive breast cancer, we then queried whether in the METABRIC cohort, there was a clear association with estrogen pathways and GADD45A expression. Using GSEA analysis, we observed that GADD45A expression was associated with several important clinical pathways (**Figure 4A**). When focusing on estrogen related pathways, we observed that GADD45A expression was associated with estrogen late response (HALLMARK_ESTROGEN_RESPONSE_LATE), with NES score of 1.23 (**Figure 4B**), suggesting a potential link between GADD45A expression and estrogen signaling.

**Figure 4.**
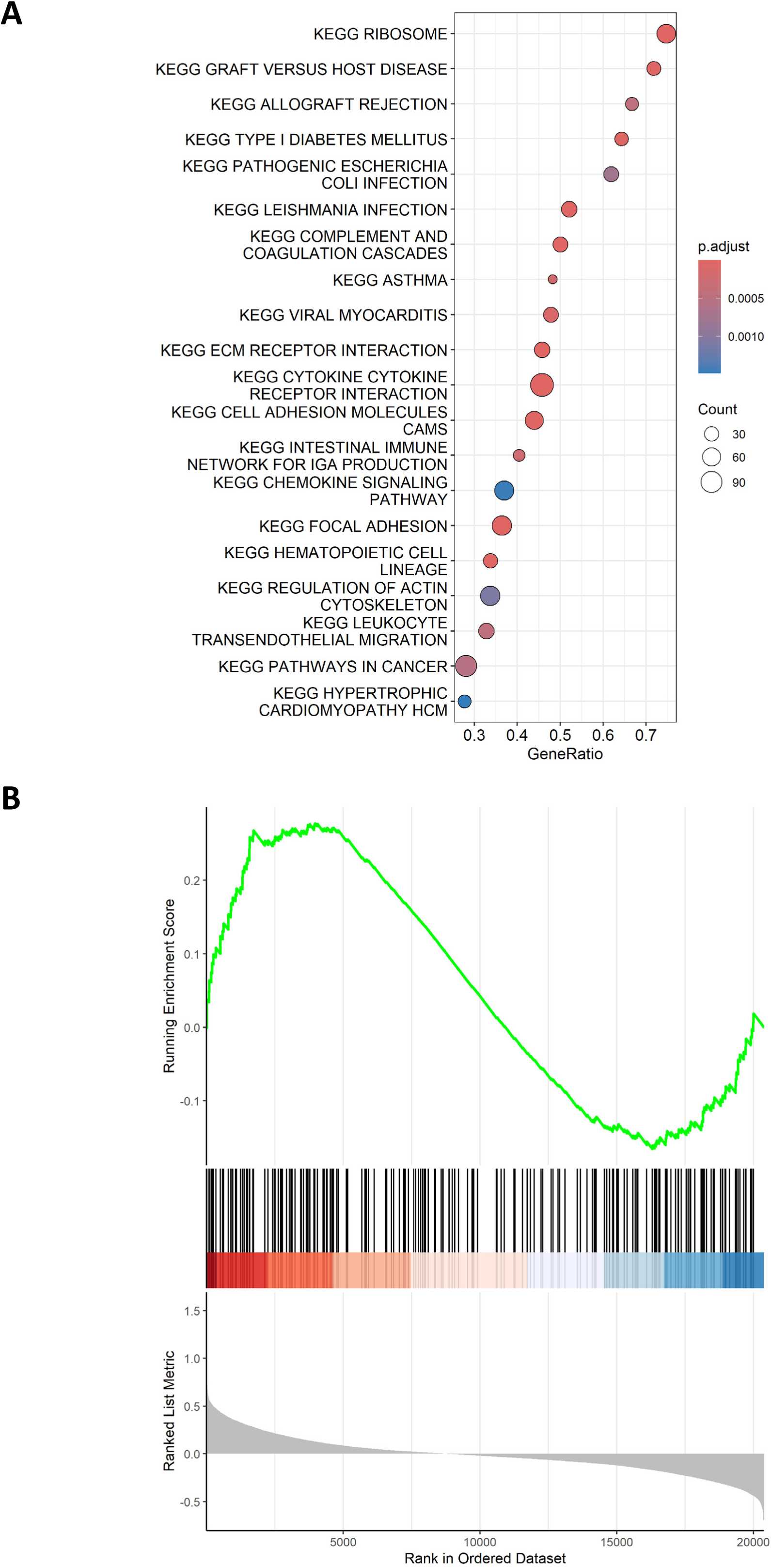
Functional analysis among the estrogen and GADD45A. (**A**), Bubble chart showing results of the GSEA analysis from samples in METABRIC cohort. (**B**), GSEA analyses of HALLMARK_ESTROGEN_RESPONSE_LATE in HR+ breast cancer from METABRIC cohort.

## Discussion

GADD45A is a nuclear protein that was originally discovered as inducible in response to DNA damage and cytotoxic therapies^18^. Its role in breast cancer was reported in a recent study where higher expression was associated with higher Ki-67 and with worse outcome in TNBC patients ^19^. Another study reported a different angle, that GADD45A was more associated with HR(+) breast cancer compared to other subtypes. Experimental work revealed the dual roles of GADD45A, where it acted as a tumor suppressor role in ras driven breast cancers and pro-tumorogenic role in myc driven breast cancer ^11^. This finding suggests that GADD45A possess multiple roles in the nucleus, where it regulates different tumor related pathways, including GSK3β/β- catenin or JNK/p38 pathways ^11, 20^. However, the exact role for GADD45A in breast cancer, especially hormone positive breast cancer, is unclarified.

Wang et al proposed a 22-gene target transcriptional signature of the Hippo pathway ^21^. The genes curated were selected from pooled RNA-seq and ChIPseq data, to identify genes that robustly alter expression levels when YAP/TAZ of the hippo pathway was inhibited ^21^. GADD45A was included in the 22-gene list. Our recent study examined RNA-seq in hormone positive breast cancer cells treated with HDAC inhibitors ^8^. Interestingly, we discovered a link between HDAC inhibitors and the hippo pathway, which we verified experimentally. When combining the finding with survival analysis, there was a strong association between upregulation of GADD45A as well as CCDC80 with improved outcomes in HR(+)HER2(-) breast cancer patients. Our study therefore proposed a axis of HDAC inhibition-Hippo pathway-GADD45A, suggesting that HDAC inhibitors or other cues that activated Hippo-TEAD transcriptional signaling which could upregulate GADD45A may be potentially beneficial for HR(+) breast cancer^8^. Much literature has probed the interplay between estrogen signaling and the hippo pathway ^22–25^. Our previous study confirmed the causal relationship between Hippo pathway and GADD45A expression^8^, although whether estrogen signaling plays a role in this relationship is unclear.

In our current study, we identify an association between higher GADD45A expression and higher ER positivity and lower Ki-67 levels. This suggests that stronger ER signaling, i.e. a “more luminal A resemblance” might be associated with higher GADD45A levels. It is unclear the exact causative relationship between estrogen and GADD45A. GADD45A is not a direct target for ER binding^26^. The observed concomitant high GADD45A expression in luminal breast cancer tumors suggest that either GADD45A regulates estrogen signaling and/or expression, or GADD45A is an indirect downstream consequence of ER signaling. The latter is compatible with a ER- Hippo-GADD45A axis, which would indicate GADD45A as a marker for estrogen signaling. It will be clinically useful and relevant to understand whether GADD45A levels respond to endocrine treatment, which would clarify a role for GADD45A as a biomarker for treatment response and efficacy assessment.

CDK4/6 inhibitors have been well established as standard of care in HR(+) HER2 (-) breast cancer in the past years. In our study, combining two independent cohorts, we did not observe an obvious relationship between PFS on CDK4/6 inhibitors and GADD45A levels, even though the samples sizes are small. Our finding is in line that GADD45A is more associated with endocrine signaling, which is less directly related to CDK4/6 inhibitor mechanisms. Thus, GADD45A levels may not directly affect therapeutic responses to CDK4/6 inhibitors. It remains unclear whether therapeutic modalities that predominantly focus on endocrine blockade, including conventional aromatase inhibitors and SERMs, ER PROTACs^27^, newer oral SERDs ^6, 28^, could change GADD45A levels, and whether GADD45A may play a more proactive role in related signaling.

In summary, our study provides further experimental data that GADD45A levels are associated with tumors that have more prominent estrogen signaling, and is associated with a better outcome in luminal cancers. Future work will further unveil the interplay between GADD45A, including its relationship with DNA damage, and the estrogen signaling network, with hopes of uncovering a more efficacious approach to target hormone receptor positive breast cancer.

## Authors contributions

Data collection, analysis and curation: Chih-Yi Lin, Jiun-I Lai, drafting of the manuscript: Chih-Yi Lin, Jiun-I Lai, conceptualization of the project: Chun-Yu Liu, Ta-Chung Chao, Chi-Cheng Huang, Yi-Fang Tsai, Ling-Ming Tseng, Jiun-I Lai. All authors read and approved the final manuscript.

## Declarations

Conflict of interest the authors declare that they have no conflict of interest.

## References

1. Siegel, R.L., Miller, K.D., Wagle, N.S. & Jemal, A. Cancer statistics, 2023. CA Cancer J Clin 73, 17–48 (2023).

2. Sung, H. et al. Global Cancer Statistics 2020: GLOBOCAN Estimates of Incidence and Mortality Worldwide for 36 Cancers in 185 Countries. CA Cancer J Clin 71, 209–249 (2021).

3. DeSantis, C.E. et al. Breast cancer statistics, 2019. CA Cancer J Clin 69, 438–451 (2019).

4. Liedtke, C. et al. Response to neoadjuvant therapy and long-term survival in patients with triple-negative breast cancer. J Clin Oncol 26, 1275–1281 (2008).

5. Spring, L.M. et al. Cyclin-dependent kinase 4 and 6 inhibitors for hormone receptor-positive breast cancer: past, present, and future. Lancet 395, 817–827 (2020).

6. Bidard, F.C. et al. Elacestrant (oral selective estrogen receptor degrader) Versus Standard Endocrine Therapy for Estrogen Receptor-Positive, Human Epidermal Growth Factor Receptor 2-Negative Advanced Breast Cancer: Results From the Randomized Phase III EMERALD Trial. J Clin Oncol 40, 3246–3256 (2022).

7. Dilawari, A. et al. US Food and Drug Administration Approval Summary: Capivasertib With Fulvestrant for Hormone Receptor-Positive, Human Epidermal Growth Factor Receptor 2-Negative Locally Advanced or Metastatic Breast Cancer With PIK3CA/AKT1/PTEN Alterations. Journal of clinical oncology: official journal of the American Society of Clinical Oncology 42, 4103–4113 (2024).

8. Lin, T.I. et al. HDAC inhibitors modulate Hippo pathway signaling in hormone positive breast cancer. Clin Epigenetics 17, 37 (2025).

9. Liebermann, D.A. & Hoffman, B. Gadd45 in stress signaling. J Mol Signal 3, 15 (2008).

10. Humayun, A. & Fornace, A.J., Jr. GADD45 in Stress Signaling, Cell Cycle Control, and Apoptosis. Adv Exp Med Biol 1360, 1–22 (2022).

11. Tront, J.S., Huang, Y., Fornace, A.J., Jr., Hoffman, B. & Liebermann, D.A. Gadd45a functions as a promoter or suppressor of breast cancer dependent on the oncogenic stress. Cancer research 70, 9671–9681 (2010).

12. Shan, Z., Li, G., Zhan, Q. & Li, D. Gadd45a inhibits cell migration and invasion by altering the global RNA expression. Cancer Biol Ther 13, 1112–1122 (2012).

13. Tront, J.S., Willis, A., Huang, Y., Hoffman, B. & Liebermann, D.A. Gadd45a levels in human breast cancer are hormone receptor dependent. Journal of translational medicine 11, 131 (2013).

14. Cerami, E. et al. The cBio cancer genomics portal: an open platform for exploring multidimensional cancer genomics data. Cancer Discov 2, 401–404 (2012).

15. Tang, Z. et al. GEPIA: a web server for cancer and normal gene expression profiling and interactive analyses. Nucleic acids research 45, W98–W102 (2017).

16. Harbeck, N. et al. Adjuvant abemaciclib combined with endocrine therapy for high-risk early breast cancer: updated efficacy and Ki-67 analysis from the monarchE study. Ann Oncol 32, 1571–1581 (2021).

17. Park, Y.H. et al. Longitudinal multi-omics study of palbociclib resistance in HR- positive/HER2-negative metastatic breast cancer. Genome Med 15, 55 (2023).

18. Rosemary Siafakas, A. & Richardson, D.R. Growth arrest and DNA damage-45 alpha (GADD45alpha). The international journal of biochemistry & cell biology 41, 986–989 (2009).

19. Wang, J. et al. The expression and clinical significance of GADD45A in breast cancer patients. PeerJ 6, e5344 (2018).

20. Tront, J.S., Hoffman, B. & Liebermann, D.A. Gadd45a suppresses Ras-driven mammary tumorigenesis by activation of c-Jun NH2-terminal kinase and p38 stress signaling resulting in apoptosis and senescence. Cancer research 66, 8448–8454 (2006).

21. Wang, Y. et al. Comprehensive Molecular Characterization of the Hippo Signaling Pathway in Cancer. Cell reports 25, 1304–1317 e1305 (2018).

22. Sadri, F., Hosseini, S.F., Rezaei, Z. & Fereidouni, M. Hippo-YAP/TAZ signaling in breast cancer: Reciprocal regulation of microRNAs and implications in precision medicine. Genes Dis 11, 760–771 (2024).

23. Lin, Q. & Yang, W. The Hippo-YAP/TAZ pathway mediates geranylgeranylation signaling in breast cancer progression. Molecular & cellular oncology 3, e969638 (2016).

24. Britschgi, A. et al. The Hippo kinases LATS1 and 2 control human breast cell fate via crosstalk with ERalpha. Nature 541, 541–545 (2017).

25. Ma, S. et al. Transcriptional repression of estrogen receptor alpha by YAP reveals the Hippo pathway as therapeutic target for ER(+) breast cancer. Nature communications 13, 1061 (2022).

26. Carroll, J.S. et al. Chromosome-wide mapping of estrogen receptor binding reveals long-range regulation requiring the forkhead protein FoxA1. Cell 122, 33–43 (2005).

27. Mario Campone, C.X.M., Michelino De Laurentiis, Hiroji Iwata, Sara A. Hurvitz, Seth Andrew Wander, Michael A. Danso, Dongrui Ray Lu, Julia Perkins Smith, Yuan Liu, Lana Tran, Sibyl Anderson, Erika P. Hamilton in ASCO, Vol. TPS1122 (2023).

28. Julia Cipriano, M. EMBER: Imlunestrant in Estrogen Receptor–Positive, HER2-Negative Advanced Breast Cancer. (2024).

