## Supplementary material for "Association of GADD45A and favorable outcome in hormone positive breast cancer": Table1, 2

#### Table1. Clinical characteristics for the tissue microarray cohort BC081116e.

| Pos. | No. | Age | Sex | Organ/Anatomic Site | Pathology diagnosis | TNM | Grade | Stage | Type | ER | PR | Ki67 | HER2 |
|---|---|---|---|---|---|---|---|---|---|---|---|---|---|
| A1 | 1 | 41 | F | Breast | Invasive carcinoma of no special type | T2N0M0 | 1 | IIA | Malignant | +++ | +++ | +,30% | 0 |
| A2 | 2 | 41 | F | Breast | Invasive carcinoma of no special type | T2N0M0 | 1 | IIA | Malignant | +++ | +++ | +,2% | 1+ |
| A3 | 3 | 40 | F | Breast | Invasive carcinoma of no special type | T2N1M0 | 2 | IIB | Malignant | + | + | +,3% | 2+ |
| A4 | 4 | 72 | F | Breast | Invasive carcinoma of no special type | T2N0M0 | 1 | IIA | Malignant | +++ | + | +,5% | 0 |
| A5 | 5 | 40 | F | Breast | Invasive carcinoma of no special type | T2N1M0 | 2 | IIB | Malignant | +++ | ++ | +,15% | 1+ |
| A6 | 6 | 44 | F | Breast | Invasive carcinoma of no special type | T2N2M0 | 2 | IIIA | Malignant | ++ | + | +,15% | 3+ |
| A7 | 7 | 57 | F | Breast | Invasive carcinoma of no special type | T4N0M0 | 2 | IIIB | Malignant | ++ | - | +,20% | 3+ |
| A8 | 8 | 46 | F | Breast | Invasive carcinoma of no special type | T2N0M0 | 2 | IIA | Malignant | - | - | +,20% | 0 |
| A9 | 9 | 27 | F | Breast | Invasive carcinoma of no special type | T3N0M0 | 2 | IIB | Malignant | + | - | +,5% | 0 |
| A10 | 10 | 59 | F | Breast | Invasive carcinoma of no special type | T2N1M0 | 2 | IIB | Malignant | - | - | +,15% | 0 |
| B1 | 11 | 41 | F | Breast | Invasive carcinoma of no special type | T3N2M0 | 2 | IIIA | Malignant | +++ | - | +,15% | 0 |
| B2 | 12 | 41 | F | Breast | Invasive carcinoma of no special type | T2N0M0 | 2 | IIA | Malignant | - | - | +,2% | 3+ |
| B3 | 13 | 48 | F | Breast | Invasive carcinoma of no special type | T3N0M0 | 2 | IIB | Malignant | - | - | - | 0 |
| B4 | 14 | 47 | F | Breast | Invasive carcinoma of no special type | T2N0M0 | 2 | IIA | Malignant | +++ | +++ | +,20% | 0 |
| B5 | 15 | 56 | F | Breast | Invasive carcinoma of no special type | T2N0M0 | 2 | IIA | Malignant | +++ | +++ | +,15% | 1+ |
| B6 | 16 | 50 | F | Breast | Invasive carcinoma of no special type | T2N0M0 | 2 | IIA | Malignant | +++ | +++ | +,10% | 0 |
| B7 | 17 | 42 | F | Breast | Invasive carcinoma of no special type | T2N2M0 | 2 | IIIA | Malignant | - | - | - | 0 |
| B8 | 18 | 54 | F | Breast | Invasive carcinoma of no special type | T2N0M0 | 2 | IIA | Malignant | - | - | +,80% | 0 |
| B9 | 19 | 64 | F | Breast | Invasive carcinoma of no special type | T2N1M0 | 2 | IIB | Malignant | +++ | +++ | +,5% | 0 |
| B10 | 20 | 47 | F | Breast | Invasive carcinoma of no special type | T2N0M0 | 2 | IIA | Malignant | +++ | - | +,15% | 0 |
| C1 | 21 | 42 | F | Breast | Invasive carcinoma of no special type | T2N0M0 | 2 | IIA | Malignant | - | - | +,5% | 3+ |
| C2 | 22 | 42 | F | Breast | Invasive carcinoma of no special type | T3N1M0 | 2 | IIIA | Malignant | + | + | - | 0 |
| C3 | 23 | 38 | F | Breast | Invasive carcinoma of no special type | T2N0M0 | 2 | IIA | Malignant | - | - | +,20% | 0 |
| C4 | 24 | 45 | F | Breast | Invasive carcinoma of no special type | T2N0M0 | 2 | IIA | Malignant | ++ | + | +,80% | 2+ |
| C5 | 25 | 45 | F | Breast | Invasive carcinoma of no special type | T2N0M0 | 2 | IIA | Malignant | - | - | +,55% | 0 |
| C6 | 26 | 45 | F | Breast | Invasive carcinoma of no special type | T2N2M0 | 2 | IIIA | Malignant | - | - | +,40% | 0 |
| C7 | 27 | 42 | F | Breast | Invasive carcinoma of no special type | T2N0M0 | 2 | IIA | Malignant | ++ | +++ | +,5% | 2+ |
| C8 | 28 | 42 | F | Breast | Invasive carcinoma of no special type | T2N0M0 | 2 | IIA | Malignant | - | - | +,45% | 3+ |
| C9 | 29 | 48 | F | Breast | Invasive carcinoma of no special type | T3N0M0 | 2 | IIB | Malignant | +++ | +++ | +,5% | 2+ |
| C10 | 30 | 56 | F | Breast | Invasive carcinoma of no special type | T2N1M0 | 2 | IIB | Malignant | - | - | +,5% | 3+ |
| D1 | 31 | 48 | F | Breast | Invasive carcinoma of no special type | T2N0M0 | 2 | IIA | Malignant | +++ | - | +,15% | 2+ |
| D2 | 32 | 42 | F | Breast | Invasive carcinoma of no special type | T2N2M0 | 2 | IIIA | Malignant | - | - | +,20% | 3+ |
| D3 | 33 | 36 | F | Breast | Invasive carcinoma of no special type | T2N0M0 | 2 | IIA | Malignant | +++ | +++ | +,30% | 1+ |
| D4 | 34 | 48 | F | Breast | Invasive carcinoma of no special type | T2N0M0 | 2 | IIA | Malignant | - | - | +,25% | 0 |
| D5 | 35 | 46 | F | Breast | Invasive carcinoma of no special type | T2N0M0 | 2 | IIA | Malignant | +++ | +++ | +,20% | 0 |
| D6 | 36 | 36 | F | Breast | Invasive carcinoma of no special type | T2N0M0 | 2 | IIA | Malignant | + | ++ | +,25% | 3+ |
| D7 | 37 | 42 | F | Breast | Invasive carcinoma of no special type | T2N1M0 | 2 | IIB | Malignant | + | +++ | +,50% | 2+ |
| D8 | 38 | 48 | F | Breast | Invasive carcinoma of no special type | T2N0M0 | 2 | IIA | Malignant | +++ | +++ | +,15% | 2+ |
| D9 | 39 | 45 | F | Breast | Invasive carcinoma of no special type | T2N0M0 | 2 | IIA | Malignant | +++ | ++ | +,3% | 3+ |
| D10 | 40 | 45 | F | Breast | Invasive carcinoma of no special type | T2N0M0 | 2 | IIA | Malignant | - | - | +,5% | 3+ |

### Table1. Clinical characteristics for the tissue microarray cohort BC081116e.

| Pos. | No. | Age | Sex | Organ/Anatomic Site | Pathology diagnosis | TNM | Grade | Stage | Type | ER | PR | Ki67 | HER2 |
|---|---|---|---|---|---|---|---|---|---|---|---|---|---|
| E1 | 41 | 30 | F | Breast | Invasive carcinoma of no special type | T2N1M0 | 2 | IIB | Malignant | + | - | +,30% | 3+ |
| E2 | 42 | 44 | F | Breast | Invasive carcinoma of no special type | T2N0M0 | 2 | IIA | Malignant | - | - | +,10% | 3+ |
| E3 | 43 | 59 | F | Breast | Invasive carcinoma of no special type | T2N0M0 | 2 | IIA | Malignant | - | - | +,5% | 3+ |
| E4 | 44 | 45 | F | Breast | Invasive carcinoma of no special type | T2N2aM0 | 2 | IIIA | Malignant | - | - | +,3% | 3+ |
| E5 | 45 | 59 | F | Breast | Invasive carcinoma of no special type | T2N0M0 | 2 | IIA | Malignant | - | - | +,15% | 0 |
| E6 | 46 | 43 | F | Breast | Invasive carcinoma of no special type | T2N0M0 | 2 | IIA | Malignant | + | - | +,40% | 0 |
| E7 | 47 | 50 | F | Breast | Invasive carcinoma of no special type | T2N0M0 | 2 | IIA | Malignant | +++ | +++ | +,15% | 0 |
| E8 | 48 | 48 | F | Breast | Invasive carcinoma of no special type | T2N2M0 | 2 | IIIA | Malignant | +++ | +++ | +,10% | 0 |
| E9 | 49 | 53 | F | Breast | Invasive carcinoma of no special type | T2N0M0 | 2 | IIA | Malignant | + | + | +,10% | 0 |
| E10 | 50 | 38 | F | Breast | Invasive carcinoma of no special type | T3N0M0 | 2 | IIB | Malignant | - | - | +,15% | 0 |
| F1 | 51 | 49 | F | Breast | Invasive carcinoma of no special type | T1N0M0 | 2 | IA | Malignant | +++ | +++ | +,5% | 0 |
| F2 | 52 | 65 | F | Breast | Invasive carcinoma of no special type | T2N0M0 | 2 | IIA | Malignant | - | - | +,5% | 3+ |
| F3 | 53 | 34 | F | Breast | Invasive carcinoma of no special type | T2N1M0 | 2 | IIB | Malignant | - | - | +,30% | 0 |
| F4 | 54 | 70 | F | Breast | Invasive carcinoma of no special type | T4N0M0 | 2 | IIIB | Malignant | - | - | +,15% | 3+ |
| F5 | 55 | 43 | F | Breast | Invasive carcinoma of no special type | T4N1M0 | 2 | IIIB | Malignant | +++ | +++ | +,45% | 1+ |
| F6 | 56 | 56 | F | Breast | Invasive carcinoma of no special type | T2N2M0 | 2 | IIIA | Malignant | +++ | ++ | +,3% | 2+ |
| F7 | 57 | 47 | F | Breast | Invasive carcinoma of no special type | T4N0M0 | 2 | IIIB | Malignant | +++ | - | +,1% | 2+ |
| F8 | 58 | 52 | F | Breast | Invasive carcinoma of no special type | T2N2M0 | 2 | IIIA | Malignant | +++ | +++ | +,3% | 1+ |
| F9 | 59 | 44 | F | Breast | Invasive carcinoma of no special type | T2N1M0 | 2 | IIB | Malignant | + | + | +,50% | 3+ |
| F10 | 60 | 46 | F | Breast | Invasive carcinoma of no special type | T2N2M0 | 2 | IIIA | Malignant | +++ | + | +,15% | 1+ |
| G1 | 61 | 39 | F | Breast | Invasive carcinoma of no special type | T2N1M0 | 2 | IIB | Malignant | +++ | +++ | +,10% | 1+ |
| G2 | 62 | 38 | F | Breast | Invasive carcinoma of no special type | T2N2M0 | 2 | IIIA | Malignant | ++ | +++ | +,10% | 0 |
| G3 | 63 | 55 | F | Breast | Invasive carcinoma of no special type | T2N0M0 | 2 | IIA | Malignant | - | - | +,15% | 0 |
| G4 | 64 | 48 | F | Breast | Breast carcinoma with apocrine differentiation | T3N1M0 | - | IIIA | Malignant | - | - | +,5% | 0 |
| G5 | 65 | 69 | F | Breast | Invasive carcinoma of no special type | T3N0M0 | 2 | IIB | Malignant | +++ | + | +,25% | 1+ |
| G6 | 66 | 56 | F | Breast | Invasive carcinoma of no special type | T2N0M0 | 3 | IIA | Malignant | +++ | +++ | +,20% | 1+ |
| G7 | 67 | 43 | F | Breast | Invasive carcinoma of no special type | T1N1M0 | 3 | IIA | Malignant | - | - | +,30% | 0 |
| G8 | 68 | 33 | F | Breast | Invasive carcinoma of no special type | T2N3M0 | 3 | IIIC | Malignant | +++ | ++ | +,5% | 3+ |
| G9 | 69 | 72 | F | Breast | Invasive carcinoma of no special type | T2N0M0 | 3 | IIA | Malignant | - | - | +,15% | 0 |
| G10 | 70 | 44 | F | Breast | Invasive carcinoma of no special type | T2N1M0 | 3 | IIB | Malignant | - | - | +,15% | 3+ |
| H1 | 71 | 40 | F | Breast | Invasive carcinoma of no special type | T2N2M0 | 2 | IIIA | Malignant | +++ | + | +,35% | 0 |
| H2 | 72 | 42 | F | Breast | Invasive carcinoma of no special type | T2N1M0 | 3 | IIB | Malignant | +++ | - | +,20% | 1+ |
| H3 | 73 | 40 | F | Breast | Invasive carcinoma of no special type | T2N0M0 | 3 | IIA | Malignant | - | - | +,30% | 0 |
| H4 | 74 | 52 | F | Breast | Invasive carcinoma of no special type | T2N0M0 | 3 | IIA | Malignant | +++ | +++ | +,25% | 0 |
| H5 | 75 | 45 | F | Breast | Invasive carcinoma of no special type | T2N2M0 | 3 | IIIA | Malignant | - | - | +,40% | 2+ |
| H6 | 76 | 42 | F | Breast | Invasive carcinoma of no special type | T2N0M0 | 3 | IIA | Malignant | - | - | +,50% | 0 |
| H7 | 77 | 33 | F | Breast | Invasive carcinoma of no special type(sparse) | T3N0M0 | 3 | IIB | Malignant | - | - | - | 3+ |
| H8 | 78 | 49 | F | Breast | Invasive carcinoma of no special type | T4N0M0 | 3 | IIIB | Malignant | + | + | +,20% | 1+ |
| H9 | 79 | 42 | F | Breast | Invasive carcinoma of no special type (spars e) | T2N0M0 | * | IIA | Malignant | ++ | ++ | +,5% | 0 |
| H10 | 80 | 37 | F | Breast | Invasive carcinoma of no special type | T3N0M0 | 3 | IIB | Malignant | + | - | +,50% | 0 |

#### Table1. Clinical characteristics for the tissue microarray cohort BC081116e.

| Pos. | No. | Age | Sex | Organ/Anatomic Site | Pathology diagnosis | TNM | Grade | Stage | Type | ER | PR | Ki67 | HER2 |
|---|---|---|---|---|---|---|---|---|---|---|---|---|---|
| I1 | 81 | 59 | F | Breast | Invasive carcinoma of no special type | T2N0M0 | 3 | IIA | Malignant | - | - | +,3% | 1+ |
| I2 | 82 | 45 | F | Breast | Invasive carcinoma of no special type | T2N0M0 | 3 | IIA | Malignant | - | - | +,35% | 0 |
| I3 | 83 | 41 | F | Breast | Invasive carcinoma of no special type (hyperp lasia of lobular tissue) | T3N1M0 | * | IIIA | Malignant | * | * | * | * |
| I4 | 84 | 39 | F | Breast | Invasive carcinoma of no special type | T2N0M0 | 3 | IIA | Malignant | +++ | +++ | +,15% | 0 |
| I5 | 85 | 44 | F | Breast | Invasive carcinoma of no special type | T3N2M0 | 3 | IIIA | Malignant | - | - | +,60% | 0 |
| I6 | 86 | 43 | F | Breast | Invasive carcinoma of no special type | T2N2aM0 | 3 | IIIA | Malignant | - | - | +,30% | 2+ |
| I7 | 87 | 51 | F | Breast | Invasive carcinoma of no special type | T2N0M0 | 3 | IIA | Malignant | - | - | +,15% | 3+ |
| I8 | 88 | 47 | F | Breast | Invasive carcinoma of no special type | T2N0M0 | 3 | IIA | Malignant | +++ | ++ | +,5% | 0 |
| I9 | 89 | 31 | F | Breast | Invasive carcinoma of no special type | T2N0M0 | 3 | IIA | Malignant | + | - | +,40% | 0 |
| I10 | 90 | 60 | F | Breast | Invasive carcinoma of no special type | T2N0M0 | 3 | IIA | Malignant | - | - | +,15% | 0 |
| J1 | 91 | 29 | F | Breast | Invasive carcinoma of no special type | T2N0M0 | 3 | IIA | Malignant | - | - | +,15% | 3+ |
| J2 | 92 | 62 | F | Breast | Invasive carcinoma of no special type | T4N0M0 | 3 | IIIB | Malignant | - | - | +,25% | 0 |
| J3 | 93 | 44 | F | Breast | Invasive carcinoma of no special type | T2N0M0 | 3 | IIA | Malignant | + | + | +,70% | 0 |
| J4 | 94 | 63 | F | Breast | Invasive carcinoma of no special type | T3N0M0 | 3 | IIB | Malignant | - | - | +,20% | 0 |
| J5 | 95 | 34 | F | Breast | Invasive carcinoma of no special type | T2N0M0 | 3 | IIA | Malignant | - | - | +,10% | 0 |
| J6 | 96 | 62 | F | Breast | Invasive carcinoma of no special type | T4N0M0 | 3 | IIIB | Malignant | - | - | +,15% | 3+ |
| J7 | 97 | 45 | F | Breast | Invasive carcinoma of no special type | T3N0M0 | 3 | IIB | Malignant | +++ | + | +,15% | 2+ |
| J8 | 98 | 70 | F | Breast | Invasive carcinoma of no special type with ne crosis | T2N0M0 | 3 | IIA | Malignant | - | - | +,1% | 0 |
| J9 | 99 | 46 | F | Breast | Invasive carcinoma of no special type | T3N1M0 | 3 | IIIA | Malignant | - | - | +,3% | 0 |
| J10 | 100 | 40 | F | Breast | Invasive carcinoma of no special type | T2N1M0 | 3 | IIB | Malignant | - | - | +,15% | 0 |
| K1 | 101 | 47 | F | Breast | Cancer adjacent breast tissue | - | - | - | AT | ++ | - | - | 0 |
| K2 | 102 | 42 | F | Breast | Cancer adjacent breast tissue of No.79 | - | - | - | AT | ++ | ++ | - | 0 |
| K3 | 103 | 20 | F | Breast | Cancer adjacent breast tissue | - | - | - | AT | ++ | ++ | + | 0 |
| K4 | 104 | 52 | F | Breast | Cancer adjacent breast tissue (fibrous tissue) | - | * | - | AT | * | * | * | * |
| K5 | 105 | 45 | F | Breast | Cancer adjacent breast tissue of No.75 | - | - | - | AT | + | ++ | - | 0 |
| K6 | 106 | 50 | F | Breast | Cancer adjacent breast tissue (fibrous tissue) | - | * | - | AT | * | * | * | * |
| K7 | 107 | 47 | F | Breast | Cancer adjacent breast tissue | - | - | - | AT | +++ | +++ | + | 0 |
| K8 | 108 | 42 | F | Breast | Cancer adjacent breast tissue of No.27 | - | - | - | AT | + | + | + | 0 |
| K9 | 109 | 38 | F | Breast | Cancer adjacent breast tissue | - | - | - | AT | ++ | ++ | + | 0 |
| K10 | 110 | 46 | F | Breast | Cancer adjacent breast tissue | - | - | - | AT | ++ | ++ | + | 0 |
| - | 0 | 42 | M | Adrenal gland | Pheochromocytoma (tissue marker) | | - | | Malignant | | | | |

Table 2. Clinical characteristics of patients at Taipei Veterans General Hospital.

| Characteristics | Number | Percentage |
|---|---|---|
| Patients, *n* | 17 | |
| Median age, y | 61 | |
| Female, % | 100 | |
| Premenopausal | 1 | 5.88 |
| ER+ (%) | 95 | |
| PR+ (%) | 47.5 | |
| HER2+ (%) | 0 | |
| Lung metastasis | 7 | 41.18 |
| Liver metastasis | 4 | 23.53 |
| Bone metastasis | 14 | 82.35 |
| Bone only | 5 | 29.41 |
| Median PFS | 364 | |
| Median OS | 784 | |
| Response | | |
| CR | 0 | 0 |
| PR | 4 | 23.53 |
| SD | 12 | 70.59 |
| PD | 1 | 5.88 |
| Progression on first line | 10 | 58.82 |
